# The diagnostic chronic lymphocytic leukaemia genome by nanopore sequencing

**DOI:** 10.1101/750059

**Authors:** Adam Burns, Daniella DiSalvo-Williams, David Bruce, Pauline Robbe, Adele Timbs, Basile Stamatopoulos, Ruth Clifford, Maria Lopopolo, Duncan Parkes, Kate Ridout, Anna Schuh

## Abstract

Chronic lymphocytic leukaemia (CLL) is characterised by considerable clinical and biological heterogeneity, with specific recurrent genomic alterations, including *TP53* mutations, deletions of chromosome 17p, and IgHV mutational status, impacting on response to chemo-immunotherapy and targeted agents. Consequently, diagnostic screening for these predictive biomarkers is recommended in both national and international clinical guidelines. Current conventional methods, including fluorescent *in-situ* hybridisation and Sanger sequencing, exhibit shortcomings in terms of cost, speed and sensitivity, and even second-generation sequencing methods encounter technical limitations imposed by short-read lengths and bio-informatics analysis. The MinION platform from Oxford Nanopore Technologies generates exceptionally long (1-100kbp) read lengths in a short period of time and at low cost, making it a good candidate for diagnostic testing. In this paper, we present a nanopore-based CLL-specific screening assay, to simultaneously screen for both *TP53* mutations and del17p13.1, as well as determining the IgHV mutation status for a single patient in one sequencing run. We sequenced 11 CLL patients and were able to generate a full diagnostic dataset for all. We identified somatic SNVs and indels in the coding region of *TP53*, and demonstrate that, following error correction of the data, we could accurately define the somatically hypermutated IgHV region in all patients. We also demonstrated the ability of the MinION platform to detect large-scale genomic deletions through low-coverage whole-genome sequencing. We conclude that nanopore sequencing has the potential to provide accurate, low-cost and rapid diagnostic information, which could be applied to other cancer types.

## Introduction

Chronic lymphocytic leukaemia (CLL) is characterised by significant clinical and biological heterogeneity, specific recurrent genetic features^1–4^ and differential response to chemoimmunotherapy^5^ and targeted agents^6,7^. Specifically, the percentage of somatic hypermutation of the immunoglobulin locus heavy chain (IgHV) caused by activation-induced cytidine deaminase (AID) activity in the germinal centres of secondary lymphoid organs has long been used to divide CLL into two distinct groups. The degree of identity of the lgHV locus with the closest germline, expressed as a percentage, correlates with different median times to first treatment, overall survival^8,9^ and also differential response to chemoimmunotherapy^10,11^. Besides, there is a strong association between chemo-immunotherapy resistance and disruption of *TP53* either through deletions of chromosome 17p (del17p13.1)^12^ and from clonal^13–15^ or sub-clonal^16^ pathogenic SNVs/Indels mutations in the *TP53* DNA-binding domain that demand sensitive detection methods. Finally, there is increasing evidence that acquired complex karyotypic abnormalities and chromothrypsis (together called genomic complexity (GC)) are associated with poor response to targeted agents such as B-cell receptor^17^ (BCR) and BCL2^18^ inhibitors. Importantly, the utility of these predictive biomarkers for clinical management decision has been emphasised in both national and international guidelines^19–21^.

However, current conventional diagnostics, i.e fluorescent *in-situ* hybridisation (FISH) to reveal del17p13.1, Sanger sequencing of the *TP53* or IgHV genes, and karyotyping of lymphocytes have significant drawbacks with respect to overall cost, speed, sensitivity and quantity and quality of input material required. The development of short-read next-generation sequencing (NGS) platforms has revolutionised our ability to determine genomic changes in cancer. However, due to the short-read length involved, there remain significant technical challenges with NGS, in particular with reference to the analysis of privately re-arranged loci, such as IgHV, that only poorly aligns to a reference genome and also with respect to GC. Single-molecule, or third-generation, sequencing, such as that offered by the MinION platform (Oxford Nanopore Technologies, Oxford, UK) enables the sequencing of non-amplified native DNA of long linear read lengths (1-100kbp). This capability, in combination with the rapid sequencing times and low initial cost and maintenance requirements of the equipment, makes an ideal candidate for patient-near testing, and an attractive DNA sequencing device for low-to-middle-income-countries^22^.

In the current study, we developed a comprehensive, nanopore-based, rapid, low-cost alternative to conventional testing for patients with CLL, generating a full diagnostic whole genome sequencing dataset on a single platform that carries the potential of diagnostic implementation in both resource-rich and resource-poor regions of the world.

## Methods

### Sample Selection and DNA Extraction

This study was conducted in accordance with the declaration of Helsinki, approved by the institutional review board (local ethics committee of John Radcliffe Hospital) and was performed using samples collected from CLL patients after written informed consent. Our study cohort was recruited from our own CLL database, and was selected by the presence of *TP53* mutations and del17p13.1. The 11 samples included both mutated (n=5) and unmutated (n=6) IgHV genes (including three cases with two rearrangements each), seven mutations in *TP53* (five missense, one stop gain and one frameshift deletion) and four del17p13.1events (table 1), previously identified through at least one of either short-read whole genome sequencing^23^ (WGS), whole transcriptome sequencing^23,24^ (WTS), targeted deep-sequencing^25^ or Sanger sequencing^24^. Genomic DNA was extracted from peripheral blood samples of the 12 untreated CLL patients using a Qiagen DNA Blood mini kit (Qiagen, Hilden, Germany).

**Table 1.** Details of patient cohort. WT = Wild type, WGS = short-read whole-genome sequencing, TSP = short-read targeted sequencing panel, M = mutated, UM = unmutated, MM = multiple mutated, MUM = multiple unmutated, MS-D = MiSeq DNA sequencing, MS-R = MiSeq RNA sequencing.

| Sample ID | TP53 |  | del(17p) |  |  |  |  | IgHV |  |  |  |
| --- | --- | --- | --- | --- | --- | --- | --- | --- | --- | --- | --- |
|  | Variant | Detection Method(s) | Type | Start | End | Size (Mb) | Detection Method(s) | Family | Status | Germline Homology (%) | Detection Method(s) |
| ADM293 | Gly193Val | TSP | WT |  |  |  |  | VH1-69 | UM | 100 | MS-D |
| ADM455 | WT | TSP | WT |  |  |  |  | VH3-21 | M | 96.7 | MS-D |
| ADM477 | Glu457Ala | TSP | cnLOH | 1 | 18,928,388 | 18.9 |  | VH3-30, VH1-30 | MUM | 98.56, 99.65 | MS-D |
| ARC603 | Arg81X | TSP | Loss | 1 | 18,928,388 | 18.9 | SNP Array | VH3-7 | UM | 100 | MS-D |
| CLL145 | Arg273His | WGS, TSP | WT |  |  |  |  | VH3-23 | M | 95.83 | MS-D, MS-R |
| CLL331 | Arg158Serfs*6 | WGS, TSP | Loss | 1 | 20,638,596 | 20.6 | WGS, SNP Array | VH3-72 | M | 96.03 | MS-D, MS-R |
| CLL345 | Lys132Gln | WGS, TSP | WT |  |  |  |  | VH3-49 | UM | 100 | MS-D, MS-R |
| CLL347 | WT | WGS, TSP | WT |  |  |  |  | VH3-9 | UM | 100 | MS-D, MS-R |
| CLL351 | WT | WGS, TSP | WT |  |  |  |  | VH3-72, VH3-13 | MM | 92.38, 91.81 | MS-D, MS-R |
| CLL371 | WT | WGS, TSP | WT |  |  |  |  | VH3-23, VH3-30 | MM | 92.36, 90.88 | MS-D, MS-R |
| CLL374 | Ser241Phe | WGS, TSP | Loss | 0 | 24,000,000 | 24.0 | WGS, SNP Array | VH3-11 | UM | 99 | MS-D |

### IgHV Library Preparation and Sequencing

The full-length IgHV locus was amplified from genomic DNA using primers based on those described by Campbell et al^26^ targeting the IGH leader (forward) and consensus JH (reverse) regions, modified to include Oxford Nanopore barcoding tails (table 2), with all eight primers pooled to a final concentration of 2μM. The final reaction contained 8μl 2μM primer pool, 12.5μl 2x Qiagen Multiplex PCR Master Mix (Qiagen, Hilden, Germany), 2.5μl 5x Qiagen Q-solution (Qiagen, Hilden, Germany) and 2μl DNA. Cycling conditions were: 95°C for 15 minutes followed by 30 cycles of 94°C for 30 seconds, 62°C for 90 seconds and 72°C for 90 seconds, with a final extension of 72°C for 10 minutes. Following clean-up with AmpureXP beads, 0.5nM of each product was used as input for the barcoding PCR protocol (SQK-LSK108, Oxford Nanopore Technologies, Oxford, UK), in a reaction mix containing 2μl ONT barcode sequence, 50μl 2x LongAmp MasterMix (New England Biolabs, Ipswich, MA, USA) to a final volume of 100μl with nuclease-free water. Cycling conditions were: 95°C for 3 minutes, 12 cycles of 95°C for 15 seconds, 62°C for 15 seconds and 65°C for 90 seconds, with a final extension time of 65°C for 10 minutes. Following clean-up with Ampure XP beads, each sample was quantified by Qubit assay and pooled for a final concentration of 1 μg. Sequencing tethers and adaptors were ligated and the library prepared for sequencing according to the SQK-LSK108 protocol (Oxford Nanopore Technologies, Oxford, UK). Sequencing was performed using a FLO-MIN107 flowcell on a MinION mk1b instrument and run for 48 hours.

**Table 2.**
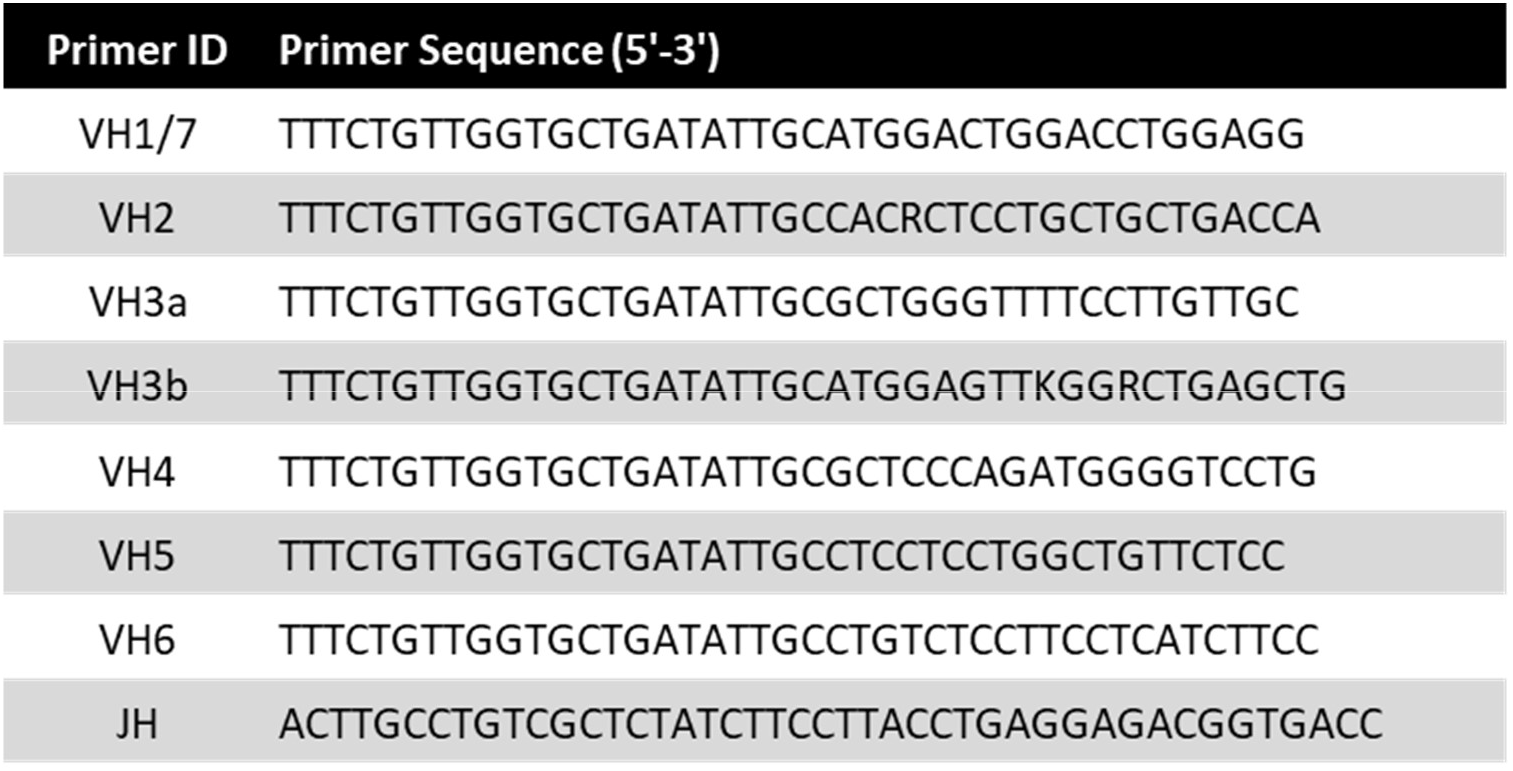
IgHV Sequencing Primers.

### TP53 Library Preparation and Sequencing

We designed primers to amplify all exons of *TP53* in 5 amplicons, with each primer including the Oxford Nanopore barcoding tail sequence (table 3). The final reaction mix contained 0.5μM forward primer, 0.5μM reverse primer, 12.5μl 2x Qiagen Multiplex Master Mix, 5μl 5x Q-solution, 3.5μl nuclease-free water and 2μl genomic DNA. For the exon 7-9 and 10-11 reactions, the Q-solution was omitted from the reaction and replaced with nuclease-free water. Cycling conditions for all amplicons were: 95°C for 15 minutes followed by 30 cycles of 94°C for 30 seconds, 62°C for 90 seconds and 72°C for 3 minutes, with a final extension of 72°C for 10 minutes. Following AMPure XP bead clean-up and Qubit quantification, each amplicon was normalised to 2nM, and combined into pools on a per-sample basis. These pools were used as input for the barcoding PCR reaction (SQK-LSK108, Oxford Nanopore Technologies, Oxford, UK), in a reaction mix containing 0.5nM amplicon pool 2μl ONT barcode sequence, 50μl 2x LongAmp MasterMix to a final volume of 100μl with nuclease-free water. Cycling conditions were: 95°C for 3 minutes, 14 cycles of 95°C for 15 seconds, 62°C for 15 seconds and 65°C for 3 minutes, with a final extension time of 65°C for 10 minutes. The resulting product was cleaned-up with AMPure XP beads, quantified with a Qubit assay and pooled for a final concentration of 1 μg. Sequencing tethers and adaptors were ligated and the library prepared for sequencing according to the SQK-LSK108 protocol (Oxford Nanopore Technologies, Oxford, UK). Sequencing was performed using a FLO-MIN107 flowcell on a MinION mk1b instrument and run for 48 hours.

**Table 3.**
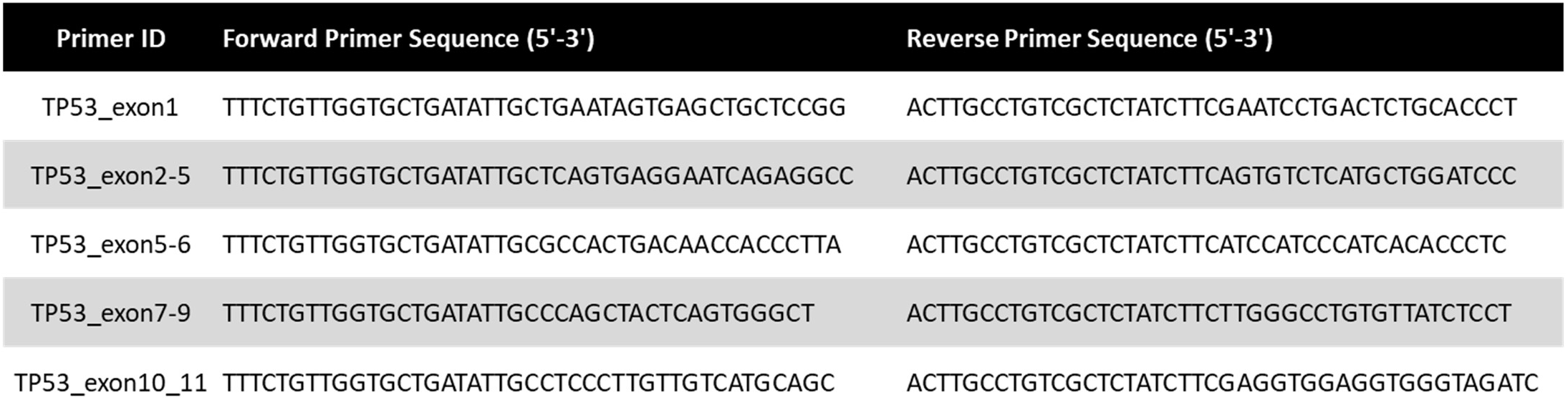
TP53 sequencing primers.

### Whole Genome Library Preparation and Sequencing

Whole genome sequencing libraries were prepared for each sample from 400ng gDNA using the Rapid Sequencing protocol (SQK-RAD004, Oxford Nanopore Technologies, Oxford, UK). Each library was sequenced independently on FLO-MIN107 flowcells, using MinION mk1b instruments with a 48-hour run time.

### Combined IgHV, TP53 and WGS assays

For the combination assay (patient CLL374), we prepared libraries for a single sample as described above. The IgHV and *TP53* libraries were assigned individual barcodes and pooled together following the barcoding PCR protocol (figure 1). A whole-genome sequencing library was prepared using the rapid sequencing kit (SQK-RAD004, Oxford Nanopore Technologies, Oxford, UK). The combination library was prepared for sequencing: 34μl sequencing buffer, 25.5μl loading beads, 2μl combined IgHV/TP53 library, 2.5μl nuclease-free water and 11 μl WGS library. The libraries were sequenced for 48 hours on R9 flowcells.

**Figure 1.**
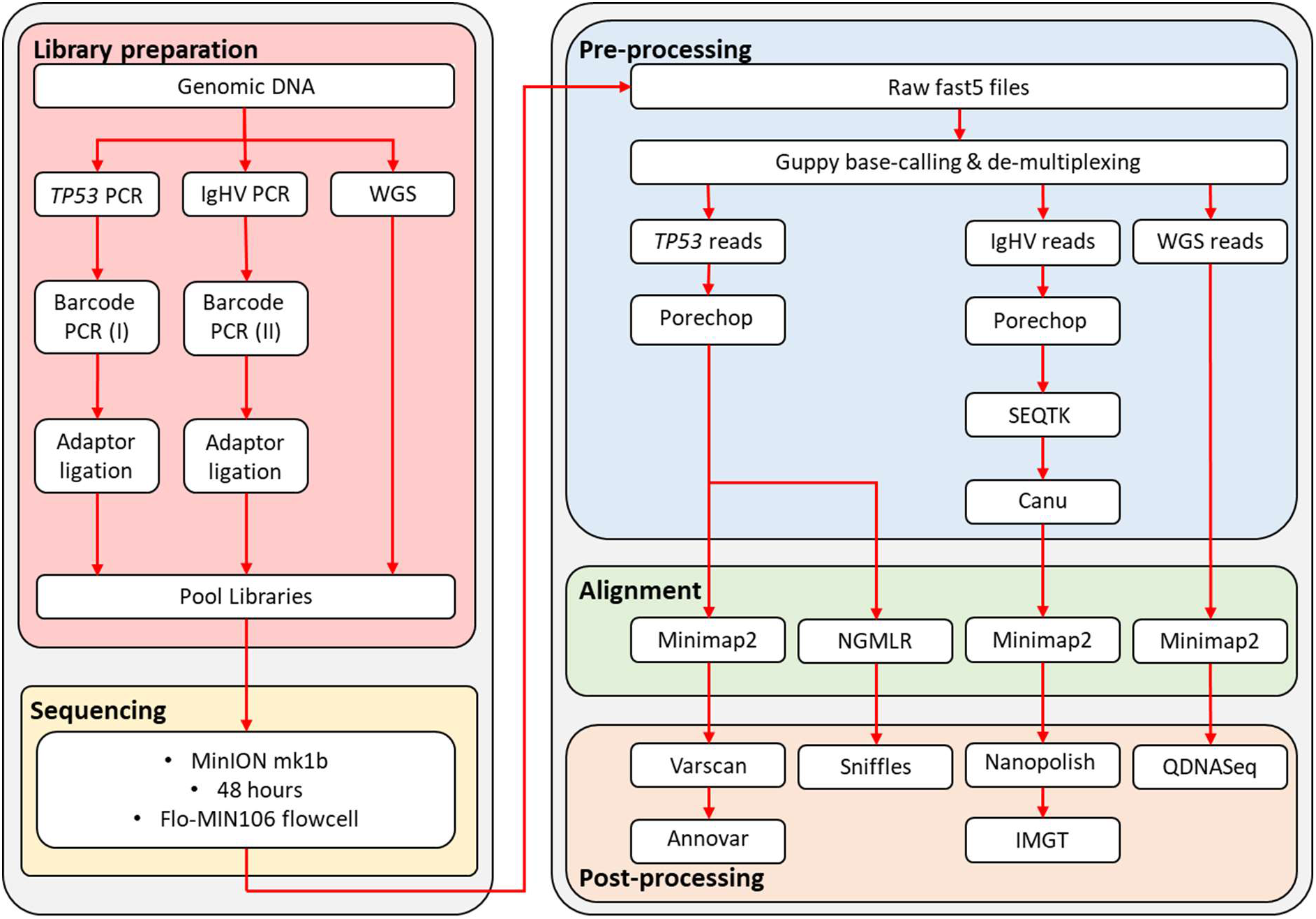
Library preparation and bio-informatic pipeline designed for IgHV sequencing, *TP53* mutation screening and copy-number analysis by nanopore sequencing in chronic lymphocytic leukaemia.

### Data Analysis

Albacore v2.1.10 (Oxford Nanopore Technologies, Oxford, UK) was used for basecalling and de-multiplexing of raw sequencing files, with Porechop (v0.2.3, https://github.com/rrwick/Porechop) used for adaptor removal (figure 1). To streamline the IgHV analysis, we down-sampled each fastq file to 10,000 reads each, using Seqtk (https://github.com/lh3/seqtk), followed by Canu^27^ to generate preliminary consensus sequences. Alignment of fastq reads, using the Canu consensus sequence as a reference, was performed with Minimap2^28^. The resulting SAM file was converted into BAM format, sorted and indexed with Samtools (v1.7, https://github.com/samtools/). A final consensus sequence was generated with Nanopolish (v0.11.0, https://github.com/its/nanopolish). The IMGT database^29,30^ was used to determine the level of identity with respect to germline, per ERIC guidelines^31^. Correlation of germline homology values was calculated using Lin’s concordance correlation coefficient (CCC) with DescTools v0.99.28 in R (version 3.5.2). For SNV detection in *TP53*, alignment files were generated by mapping reads to the hg19 genome build with Minimap2^28^. We used Samtools (v1.7, https://github.com/samtools/) to compile a pileup file using for each sample, which was used as input for VarScan^32^ v2.3.9 for variant calling. To exclude random errors and reduce false positive, the following filtering criteria were used: minimum coverage 400x, minimum VAF 0.1, minimum average quality 20. All remaining variants were annotated with Annovar^33^. To identify indels, we used NGMLR^34^, with ONT-specific options, to align our reads to hg19. Indels were called using Sniffles^34^, and only retained if identified as ‘precise’ variants by the software. Whole-genome datasets were aligned to hg19 with Minimap2^28^, and sorted and indexed with Samtools (v1.7, https://github.com/samtools/). For copy-number analysis, we used the QDNAseq^35^ package version 1.20.0 and R version 3.5.2. Briefly, aligned reads for each shallow whole-genome dataset are grouped into 100kb windows. The number of reads per window are counted, adjusted for GC content and mappability, and filtered to remove low quality regions, allowing regions of copy-number alteration to be called.

For the combined library sequencing run, reads were base-called and de-multiplexed using Guppy v3.2.1. The IgHV and *TP53* reads were split into two different barcode folders, and the remaining WGS reads grouped into the ‘unknown’ folder. Downstream analysis of these data were as described above.

## Results

### Identification of somatic mutations in TP53

We achieved an average sequencing depth of 17,087x (range 1,882-45,443) across the exonic region of *TP53*, with a raw error rate of 12.5%. After filtering the called variants for both quality and minimum coverage, we identified 46 SNVs and 21 indels across 11 samples. After eliminating variants in intronic and non-coding regions, seven SNVs remained within the coding region of *TP53* transcript NM_000546, five missense (Lys132Gln, Ser241Phe, Arg273His, Glu285Lys, Glu325Val) and 1 nonsense (Arg213X) (table 4). A 19bp frameshift deletion in exon 5 was also identified (Glu154Serfs*6) in CLL331. Generally, we observed good correlation of the variant allele frequencies between MinION and MiSeq datasets, with the exception of the Arg213X variant in ARC603 (median VAF difference between the 2 methods 0.050 (range 0.010-0.280) including Arg213X, 0.045 (range 0.010-0.110) excluding Arg213X). Closer inspection shows the Arg213 locus was only sequenced in the reverse read of the MiSeq dataset, with the substitution located at the end of an adenine tetramer (figure 2).

**Figure 2.**
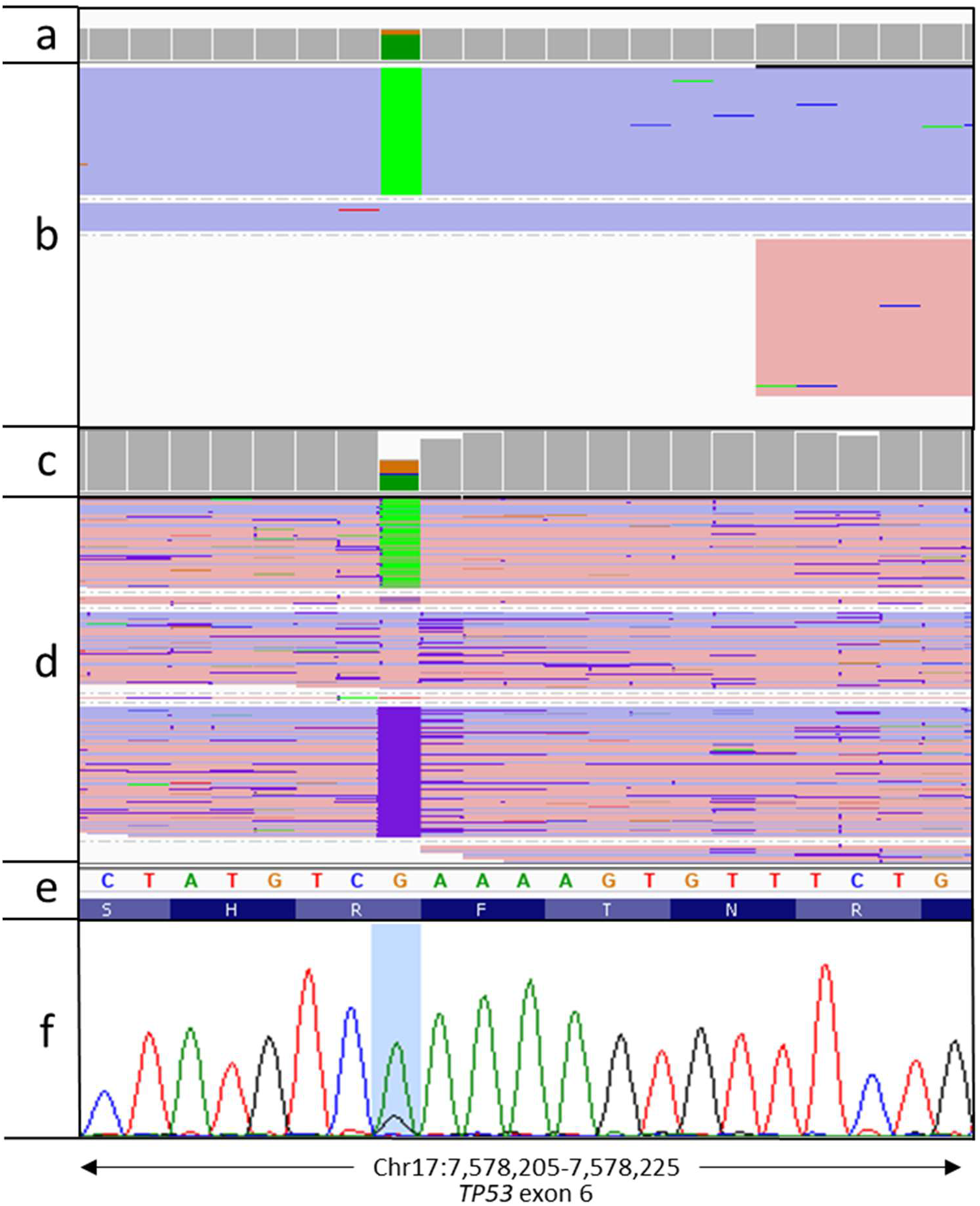
Visualisation of Arg213X variant in ARC603. **a)** sequencing read depth for each nucleotide across the region from a targeted MiSeq sequencing assay, **b)** aligned MiSeq reads with G>A variants highlighted in green. Reads are coloured according to which strand they map to (blue = positive strand, red = negative strand), **c)** MinION sequencing read depth, **d)** aligned MinION reads, **e)** genomic sequence from the hg19 genome build, **f)** confirmatory Sanger sequencing plot, with the Arg213X variant highlighted.

**Table 4.**
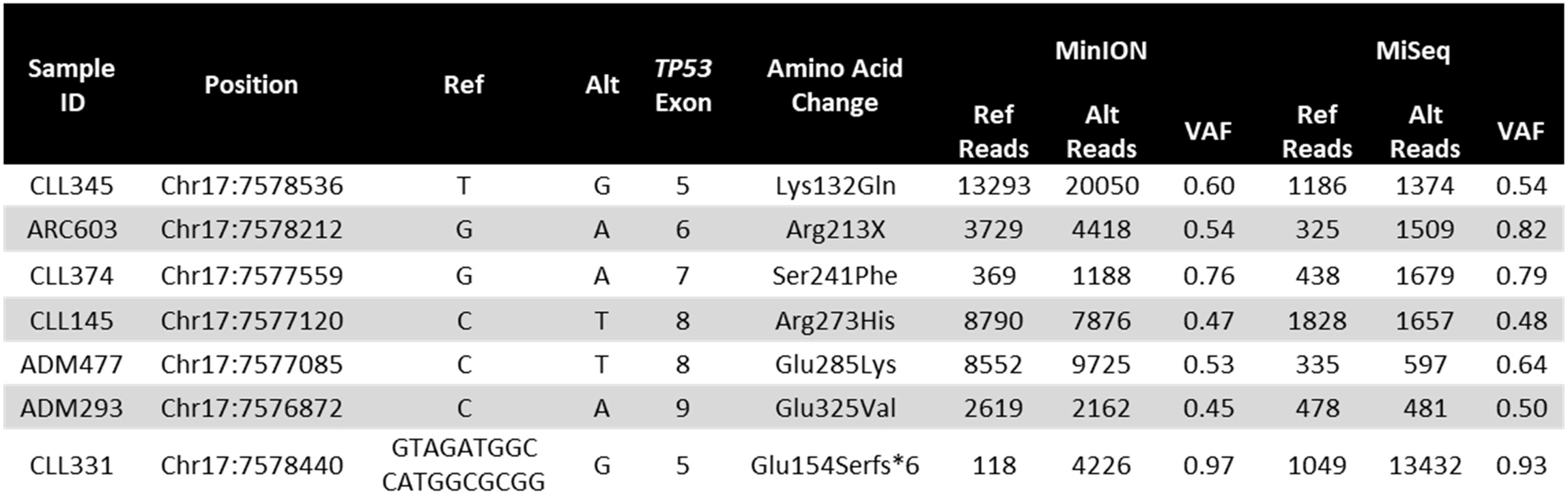
Details of variants detected in *TP53*.

### Full-length sequencing and identification of IgHV family and germline homology

Sequencing of the full-length IgHV locus for each sample generated an average of 172,747 reads (range 41,517-1,060,464). These reads were down-sampled to 10,000x for downstream analysis. In the absence of any error-correction, it is possible to identify the correct VH family for each sample, however, due to the high error rate inherent in nanopore sequencing data, it was not possible to differentiate between IgHV^mut^ and IgHV^unmut^ cases from the raw dataset. Following error correction and consensus building with Canu the germline identity improved to a level in-line with that obtained from both DNA and RNA-based short-read sequencing assays (CCC=0.998 [0.993,0.999]), although the rearrangements were deemed to be unproductive in five cases. Using Nanopolish to calculate an improved consensus sequence maintained the percentage homology information, and called a productive rearrangement in all cases (figure 3, table 5). Secondary IgH re-arrangements were accurately identified in the final data of two of the three samples previously identified (CLL371 and ADM477). The secondary IgH rearrangement in CLL351 was found in the raw nanopore data, however insufficient supporting reads resulted in it being omitted during downstream processing.

**Figure 3.**
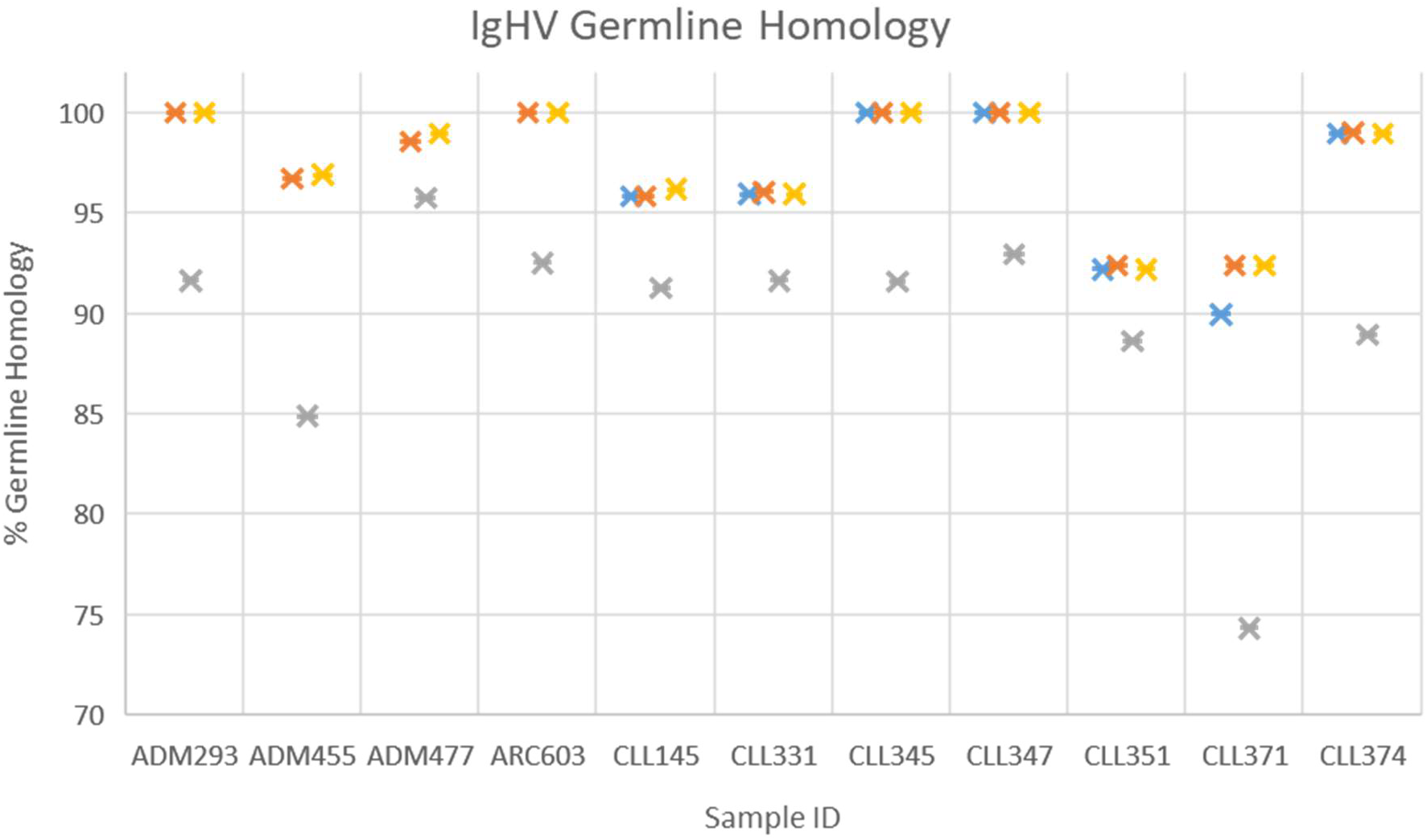
Comparison of % germline identity as determined by different sequencing techniques. Error correction of the nanopore sequencing reads is required to accurately define the germline identity of each sample. Blue = MiSeq (RNA), orange = MiSeq (DNA), grey = Raw MinION (DNA), yellow = Corrected MinION (DNA).

**Table 5.**
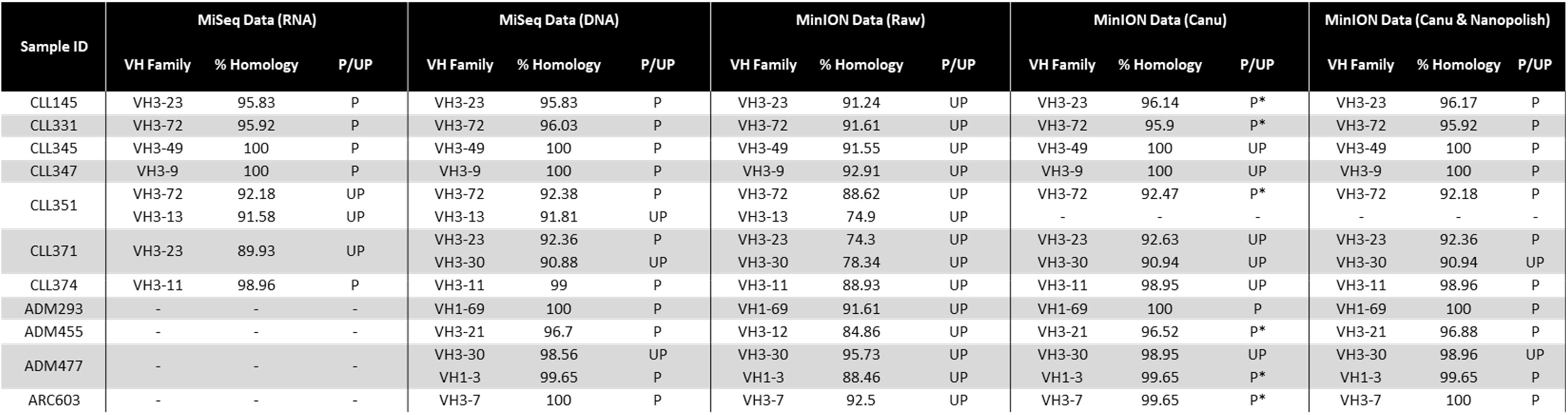
Comparison of the IgHV analysis between short-read sequencing (DNA and RNA), and nanopore sequencing before and after polishing of the data. VH Family = immunoglobulin heavy variable family, P = productive rearrangement, UP = unproductive rearrangement.

| Sample ID | MiSeq Data (RNA) |  |  | MiSeq Data (DNA) |  |  | MinION Data (Raw) |  |  | MinION Data (Canu) |  |  | MinION Data (Canu & Nanopolish) |  |  |
| --- | --- | --- | --- | --- | --- | --- | --- | --- | --- | --- | --- | --- | --- | --- | --- |
|  | VH Family | % Homology | P/UP | VH Family | % Homology | P/UP | VH Family | % Homology | P/UP | VH Family | % Homology | P/UP | VH Family | % Homology | P/UP |
| CLL145 | VH3-23 | 95.83 | P | VH3-23 | 95.83 | P | VH3-23 | 91.24 | UP | VH3-23 | 96.14 | P* | VH3-23 | 96.17 | P |
| CLL331 | VH3-72 | 95.92 | P | VH3-72 | 96.03 | P | VH3-72 | 91.61 | UP | VH3-72 | 95.9 | P* | VH3-72 | 95.92 | P |
| CLL345 | VH3-49 | 100 | P | VH3-49 | 100 | P | VH3-49 | 91.55 | UP | VH3-49 | 100 | UP | VH3-49 | 100 | P |
| CLL347 | VH3-9 | 100 | P | VH3-9 | 100 | P | VH3-9 | 92.91 | UP | VH3-9 | 100 | UP | VH3-9 | 100 | P |
| CLL351 | VH3-72 | 92.18 | UP | VH3-72 | 92.38 | P | VH3-72 | 88.62 | UP | VH3-72 | 92.47 | P* | VH3-72 | 92.18 | P |
|  | VH3-13 | 91.58 | UP | VH3-13 | 91.81 | UP | VH3-13 | 74.9 | UP | - | - | - | - | - | - |
| CLL371 | VH3-23 | 89.93 | UP | VH3-23 | 92.36 | P | VH3-23 | 74.3 | UP | VH3-23 | 92.63 | UP | VH3-23 | 92.36 | P |
|  |  |  |  | VH3-30 | 90.88 | UP | VH3-30 | 78.34 | UP | VH3-30 | 90.94 | UP | VH3-30 | 90.94 | UP |
| CLL374 | VH3-11 | 98.96 | P | VH3-11 | 99 | P | VH3-11 | 88.93 | UP | VH3-11 | 98.95 | UP | VH3-11 | 98.96 | P |
| ADM293 | - | - | - | VH1-69 | 100 | P | VH1-69 | 91.61 | UP | VH1-69 | 100 | P | VH1-69 | 100 | P |
| ADM455 | - | - | - | VH3-21 | 96.7 | P | VH3-12 | 84.86 | UP | VH3-21 | 96.52 | P* | VH3-21 | 96.88 | P |
| ADM477 | - | - | - | VH3-30 | 98.56 | UP | VH3-30 | 95.73 | UP | VH3-30 | 98.95 | UP | VH3-30 | 98.96 | UP |
|  |  |  |  | VH1-3 | 99.65 | P | VH1-3 | 88.46 | UP | VH1-3 | 99.65 | P* | VH1-3 | 99.65 | P |
| ARC603 | - | - | - | VH3-7 | 100 | P | VH3-7 | 92.5 | UP | VH3-7 | 99.65 | P* | VH3-7 | 100 | P |

### Detection of large-scale copy-number alterations

We generated a whole genome dataset with an average depth of coverage of 1.6x (range 0.1-3.4x) across all 11 samples, with an average read length of 6.3kbp (figure 4). Three samples (ARC603, CLL331 and CLL374) harboured del17p13.1, of 21.3Mb, 20.3Mb and 22Mb length, respectively (figure 5 and table 6). This matched well with both WGS and array data obtained previously. Furthermore, del(13q) was identified in four samples, with an 800kbp minimally deleted region (MDR) encompassing *MIR16, MIR15A* and *DLEU1*.

**Figure 4.**
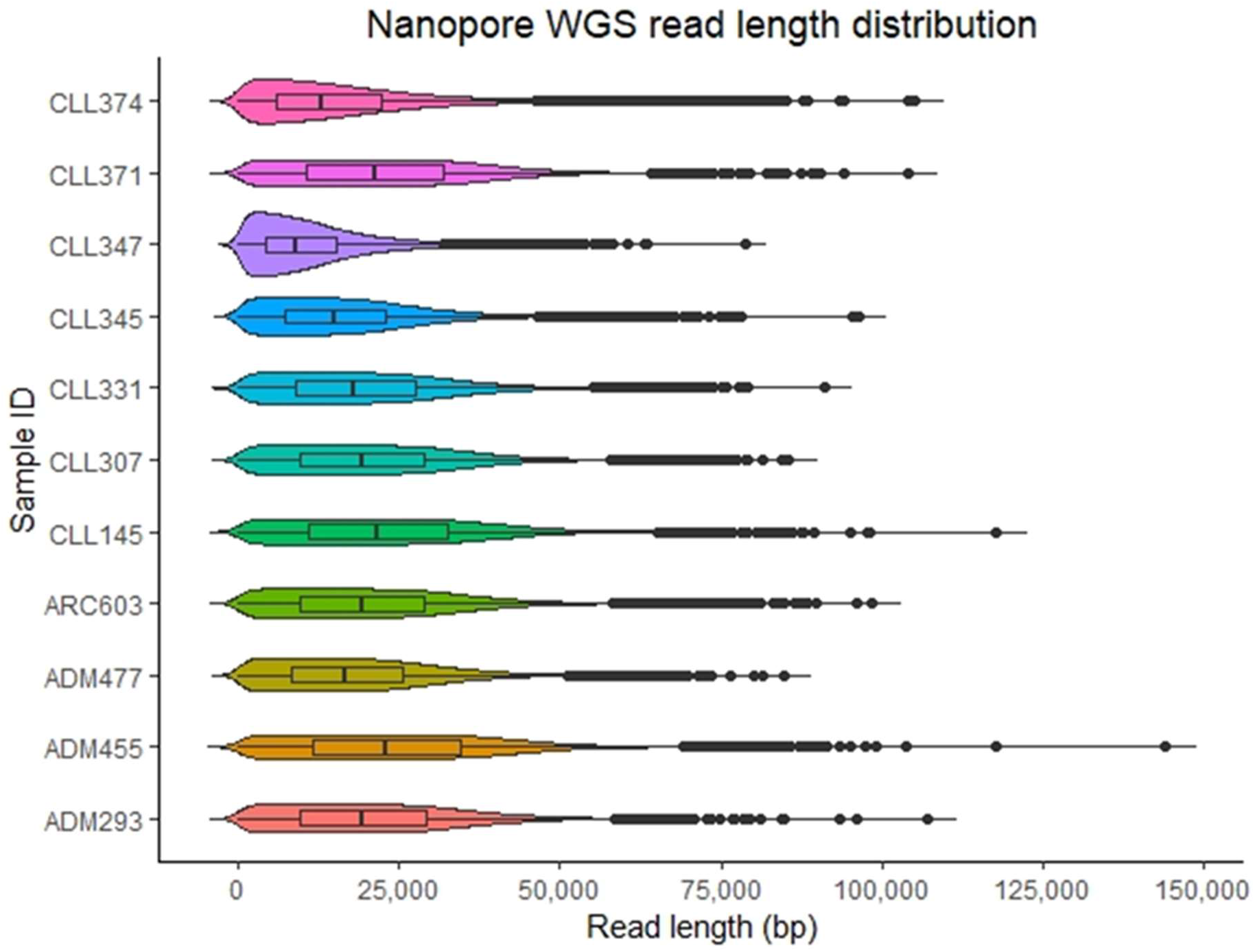
Read length distribution from MinION low-coverage whole-genome sequencing dataset.

**Figure 5.**
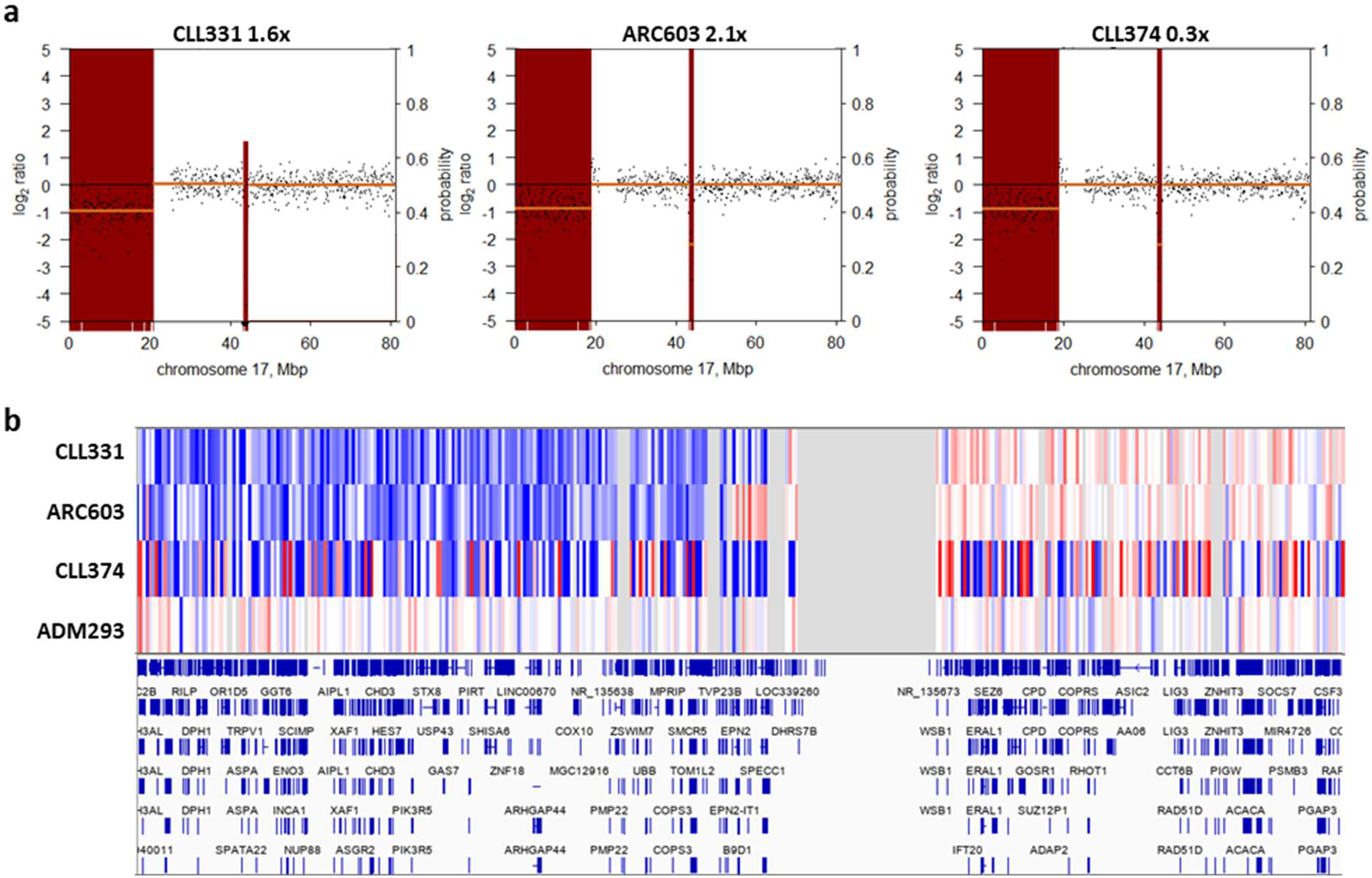
Visualisation of del(17p) in three CLL cases. a) Log2 ratio plots showing copy number profile from QDNASeq. X-axis = genomic location on chr17, primary Y-axis = log2 transformed read depth, secondary Y-axis = copy-number alteration call confidence value. b) Copy-number data for three samples with del(17p) (CLL331, ARC603 and CLL374), alongside a wild-type sample (ADM293) plotted in IGV, showing RefSeq annotation of the affected region of chr17. Blue represents regions of copy-loss.

**Table 6.**
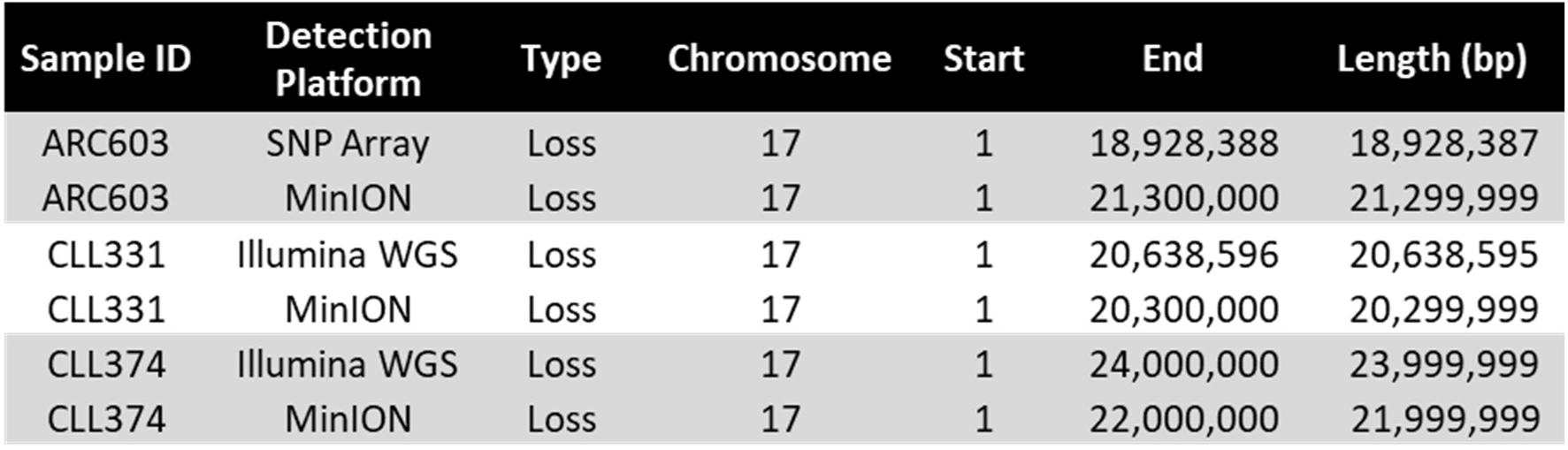
Details of the copy-number alterations detected by shallow whole-genome nanopore sequencing.

## Discussion

In summary, our data demonstrates the feasibility of nanopore sequencing as an alternative to existing, short-read, sequencing systems as a diagnostic platform. Conventional WGS remains expensive, with a recent health-economic evaluation estimating the overall costs at £6,841^36^. This is largely due to the high cost of consumables, equipment purchase and maintenance costs. We were able to generate a comprehensive suite of diagnostic information from a single sequencing run, including both targeted deep sequencing and WGS data, in ~5 days, using a platform that is both cheap to purchase and maintain, for less than £1,000. Whereas previous nanopore sequencing studies in CLL focussed only on the detection of SNVs^37^, we were able to identify both SNVs and indels, as well as accurately defining the IgHV region, a patient-specific locus subject to somatic hypermutation. The unrestricted read length of nanopore sequencing enabled us to sequence the full exonic region of *TP53*, along with full length IgHV genes, in accordance with the current ERIC guidelines for screening CLL^31,38^.

We conclude that nanopore sequencing has the potential to provide accurate, low-cost and rapid diagnostic information in a patient-near setting, and which could be applied to other cancer types.

